# Regardless of the deposition pathway, aminoacid 31 in histone variant H3 is essential at gastrulation in Xenopus

**DOI:** 10.1101/612515

**Authors:** David Sitbon, Ekaterina Boyarchuk, Geneviève Almouzni

**Affiliations:** Institut Curie, PSL Research University, CNRS, UMR3664, Equipe Labellisée Ligue contre le Cancer, 75005, Paris, France; Sorbonne Universités, UPMC Univ Paris 06, CNRS, UMR3664, 75005, Paris, France

## Abstract

The closely related replicative H3 and non-replicative H3.3 variants show specific requirement during development in vertebrates. Whether it involves distinct mode of deposition or unique roles once incorporated into chromatin remains unclear. To disentangle the two aspects, we took advantage of the Xenopus early development combined with chromatin assays. Our previous work showed that in Xenopus, depletion of the non-replicative variant H3.3 impairs development at gastrulation, without compensation through provision of the replicative variant H3.2. We systematically mutated H3.3 at each four residues that differ from H3.2 and tested their ability to rescue developmental defects. Surprisingly, all H3.3 mutated variants functionally complemented endogenous H3.3, regardless of their incorporation pathways, except for one residue. This particular residue, the serine at position 31 in H3.3, gets phosphorylated onto chromatin in a cell cycle dependent manner. While the alanine substitution failed to rescue H3.3 depletion, a phosphomimic residue sufficed. We conclude that the time of gastrulation reveals a critical importance of the H3.3S31 residue independently of the variant incorporation pathway. We discuss how this single evolutionary conserved residue conveys a unique property for this variant in vertebrates during cell cycle and cell fate commitment.

## Introduction

The organization of DNA into chromatin provides not only a means for compaction but also a versatile landscape contributing to cell fate and plasticity (Schneider and Grosschedl 2007; Yadav et al. 2018). The basic unit, the nucleosome core particle, is composed of a histone tetramer (H3-H4)_2_ flanked by two histone dimers H2A-H2B, around which 147bp of DNA are wrapped as shown by crystal structure (Luger et al. 1997). Importantly, modulation of this unit exploits the choice of histone variants and reversible PTMs (Post-translational modification) to impact cell functions (Kouzarides 2007; Gurard-Levin and Almouzni 2014). Three out of the four-histone families possess histone variants (Franklin and Zweidler 1977; Sarma and Reinberg 2005; Buschbeck and Hake 2017; Sitbon et al. 2017; Talbert and Henikoff 2017). In the H3 family, while the centromere-specific histone CENP-A (Centromere protein A) is very distinct and marks the centromere, the other well-characterized H3 histone variants are closely related and thought to ensure similar functions (Maze et al. 2014). Indeed, while they show similar structural features at the core particle level (Tachiwana et al. 2011), an important difference concerned their cell cycle regulation and distinct mode of incorporation (Mendiratta et al. 2018). In human, the two replicative variants H3.1 and H3.2 are incorporated into chromatin in a DSC (DNA-synthesis coupled) manner via the histone chaperone complex CAF-1 (Chromatin assembly factor 1) (Stillman 1986; Smith and Stillman 1989; Gaillard et al. 1996; Tagami et al. 2004; Polo et al. 2006; Groth et al. 2007; Latreille et al. 2014). By contrast, the non-replicative form H3.3, which differs from H3.1 and H3.2 by five and four residues respectively, is incorporated in a DSI (DNA-synthesis independent) manner (Ahmad and Henikoff 2002b; Ray-Gallet et al. 2002; Tagami et al. 2004; Ray-Gallet et al. 2011). This incorporation depends on the histone chaperone complex HIRA (Histone regulator A) in euchromatin regions (Lamour et al. 1995; Ray-Gallet et al. 2002; Tagami et al. 2004; Goldberg et al. 2010; Ray-Gallet et al. 2011), while the presence of H3.3 at telomeres and pericentric heterochromatin rather relies on the histone chaperone complex DAXX (Death domain-associated protein)/ATRX (Alpha thalassemia/mental retardation syndrome X-linked) (Gibbons et al. 1995; Yang et al. 1997; Drane et al. 2010; Goldberg et al. 2010; Lewis et al. 2010). Thus, the dynamics of the different histone variants with their deposition is critically linked to dedicated histone chaperones (Gurard-Levin et al. 2014; Hammond et al. 2017). Inspired by the quote “nothing in biology makes sense except in the light of evolution” (Dobzhansky 1973), we considered H3 variants in light of their conservation in different organisms. In *Saccharomyces cerevisiae*, the only non-centromeric histone H3 is mostly related to H3.3, and achieves both replicative and non-replicative variant functions, illustrating the capacity of survival with a single variant (Dion et al. 2007; Jamai et al. 2007; Rufiange et al. 2007). Intriguingly, however, humanized *S. cerevisiae* with all histones replaced by human orthologs survived better with hH3.1 than hH3.3 (Truong and Boeke 2017). The better adaptation to H3.1 could argue for non-essential roles of H3.3 in *S. cerevisae*, a unicellular organism. However, in metazoans like *Drosophila melanogaster*, the replicative variant can compensate for the loss of H3.3 during development in somatic tissues, although the adults are sterile (Loppin et al. 2005; Bonnefoy et al. 2007; Hodl and Basler 2009; Sakai et al. 2009; Orsi et al. 2013). Since the sterility could simply reflect a shortage of maternal H3.3 to replace protamine from sperm chromatin after fertilization, the most parsimonious hypothesis suggests that the nature of the variant itself might not necessarily be essential. Similarly, H3.3 is dispensable in *Caenorhabditis elegans*, since its removal does not result in lethality but rather to reduced fertility and viability in response to stress (Delaney et al. 2018). In *Arabidopsis thaliana*, in contrast, replicative and non-replicative H3 variants are clearly essential. The absence of H3.3 leads to embryonic lethality and also partial sterility due to defective male gametogenesis (Wollmann et al. 2017). In vertebrate models like *Mus musculus*, the deletion of one of the two copies of the H3.3 gene leads not only to sterility but also to developmental defects at E12.5 (Couldrey et al. 1999; Bush et al. 2013; Tang et al. 2015). Finally in human, while a complete absence is not reported, dominant effects with substitutions in H3.3, like H3.3 K27M and H3.3 G34R/V, along with mutations affecting their chaperones like DAXX/ATRX, have been involved in distinct aggressive cancers (Heaphy et al. 2011; Jiao et al. 2011; Schwartzentruber et al. 2012; Wu et al. 2012; Behjati et al. 2013; Lewis et al. 2013; Lindroth and Plass 2013; Fontebasso et al. 2014; Taylor et al. 2014; Wu et al. 2014). Furthermore, not only mutation but also expression of histone chaperones is often found altered in aggressive cancer (Corpet et al. 2011; Montes de Oca et al. 2015; Gurard-Levin et al. 2016). Thus, the developmental defects, along with situation encountered in particular cancers, underline the importance of each H3 variant and their chaperones in vertebrates. Given the limited sequence difference between H3 variants, a major puzzle is whether the need for a particular histone variant could reflect either (i) a unique mode of incorporation and provision or (ii) a distinct identity once incorporated into chromatin to drive their functions. While, the first hypothesis had been largely favored, based on previous work including ours, the issue has never been formally addressed. To disentangle these two possibilities, we decided to use *Xenopus laevis* as it is an ideal model system to tackle this issue. Indeed, extensively characterized both in developmental biology and chromatin studies (Laskey et al. 1977; Almouzni and Mechali 1988; Almouzni et al. 1990; Almouzni and Wolffe 1993), its external development permits direct access to embryos for observation and manipulation (Nieuwkoop and Faber 1994). With retention of H3 variants in sperm (Katagiri and Ohsumi 1994) and only one replicative histone H3.2, it offers a simpler situation while retaining amino acid sequence conservation with human variants for both H3.2 and H3.3. Furthermore, following fertilization, development starts by twelve rapid embryonic cell divisions, which include only S and M phases (Newport and Kirschner 1982; Etkin 1988; Masui and Wang 1998). At the MBT (midblastula transition), zygotic activation occurs concomitantly with a progressive lengthening of the cell cycle, to reach, at gastrulation, a typical cell cycle with two gap phases. In addition, cells begin to engage into defining different types with acquisition of migration properties. Importantly, previous work in our laboratory revealed a specific requirement of H3.3 during *Xenopus laevis* early development at the time of gastrulation (Szenker et al. 2012). This work showed that depletion of endogenous H3.3 (further referred to as H3.3) leads to severe gastrulation defects that cannot be rescued by providing the replicative counterpart H3.2 but only by H3.3 itself. Interestingly, there are only four residues that diverge between these two forms. A first region of divergence now known as the AIG motif, in the globular domain of H3.3, accounts for histone variant recognition by dedicated histone chaperones (Ahmad and Henikoff 2002a; Elsasser et al. 2012; Liu et al. 2012; Ricketts et al. 2015). This first region is thus key for the choice of deposition pathways. The other distinctive residue, a serine only present in H3.3, is located on the histone N-terminus, at position 31, and can be phosphorylated (Hake et al. 2005; Wong et al. 2009; Hinchcliffe et al. 2016). In our study, we systematically mutated the H3.3 histone variant in its distinct residues to assess their ability to rescue the gastrulation defects and examined their incorporation mode. In our assays, we find that mutations affecting the incorporation pathway proved neutral to set specific H3.3 functions at the time of gastrulation. By contrast, the serine at position 31 is critical in the rescue experiments after H3.3 depletion. In Xenopus, this residue, which is notably conserved in human cells, gets phosphorylated during the cell cycle with a peak in mitosis. Remarkably, in our rescue experiments, the phosphomimic form, which cannot be dynamically modified, fully rescued. We discuss how this residue and the charge at this site can contribute to establish particular chromatin states important in the context of somatic cell cycle and development.

## Results

### Conservation of histone H3 variants and their chaperones to assay H3.3 defects at gastrulation in Xenopus

While *Homo sapiens* presents two replicative H3 variants, H3.1 and H3.2, *Xenopus laevis* only possesses one replicative variant, H3.2. Both H3.2 and H3.3 are conserved with their human orthologs (Figure 1 A). Interestingly, the two H3 variants are almost identical and conserved through evolution (Waterborg 2012) (Figure 1 B). Two regions show differences in H3.2 and H3.3. The first one encompasses positions 87, 89 and 90 with a serine, a valine, and a methionine, known as the SVM motif in H3.2. Instead, these positions correspond to an alanine, an isoleucine and a glycine, known as the AIG motif in H3.3. The second difference lies at position 31 where H3.2 shows an alanine and H3.3 a serine. Considering sequences for H3.2 and H3.3 histone variants from five different model organisms in which the function along with the deposition pathways of H3 variants have been studied, the replicative variant H3.2 shows around 3% (four variable residues out of 136). In the case of the non-replicative variant H3.3, it varies of 4% differences, with six variable residues. Remarkably, the region responsible for histone chaperone recognition shows the highest variations, in line with possible coevolution with their respective histone chaperones that are not closely conserved (Supplemental figure 1 A). To examine how deposition pathways and the role for histone variants can be related, targeting these regions could thus be considered. Using *Xenopus laevis* embryos, we had previously found that a morpholino against H3.3 leads to defects during late gastrulation (Szenker et al. 2012) (Figure 1 C and Supplemental figure 1 B and Film 1). While the blastopore forms and invaginates during gastrulation, depletion of H3.3 leads to an arrest of the blastopore closure. Exogenous HA-tagged H3.3, but not HA-tagged H3.2 mRNAs (further referred as eH3s) when co-injected with the morpholino, can rescue the phenotype (Szenker et al. 2012). To optimize the assay conditions, we decided to titrate the concentration of morpholino (Supplemental figure 1 C). The lowest concentration enabled the blastopore to start to invaginate without complete closure and lead to late gastrulation defects, consistent with our previous findings (Szenker et al. 2012). By increasing by two to fourfold morpholino concentrations, gastrulation defects coincide with an even earlier phenotype, where the blastopore closure does not happen at all. These data are in line with a titration effect, whereby gastrulation time allows to readily reveal H3.3 requirements. We could thus use it as a read out to assess the role for a distinct mode of incorporation for H3 variants or a need of the histone variant itself once incorporated.

**Figure 1:**
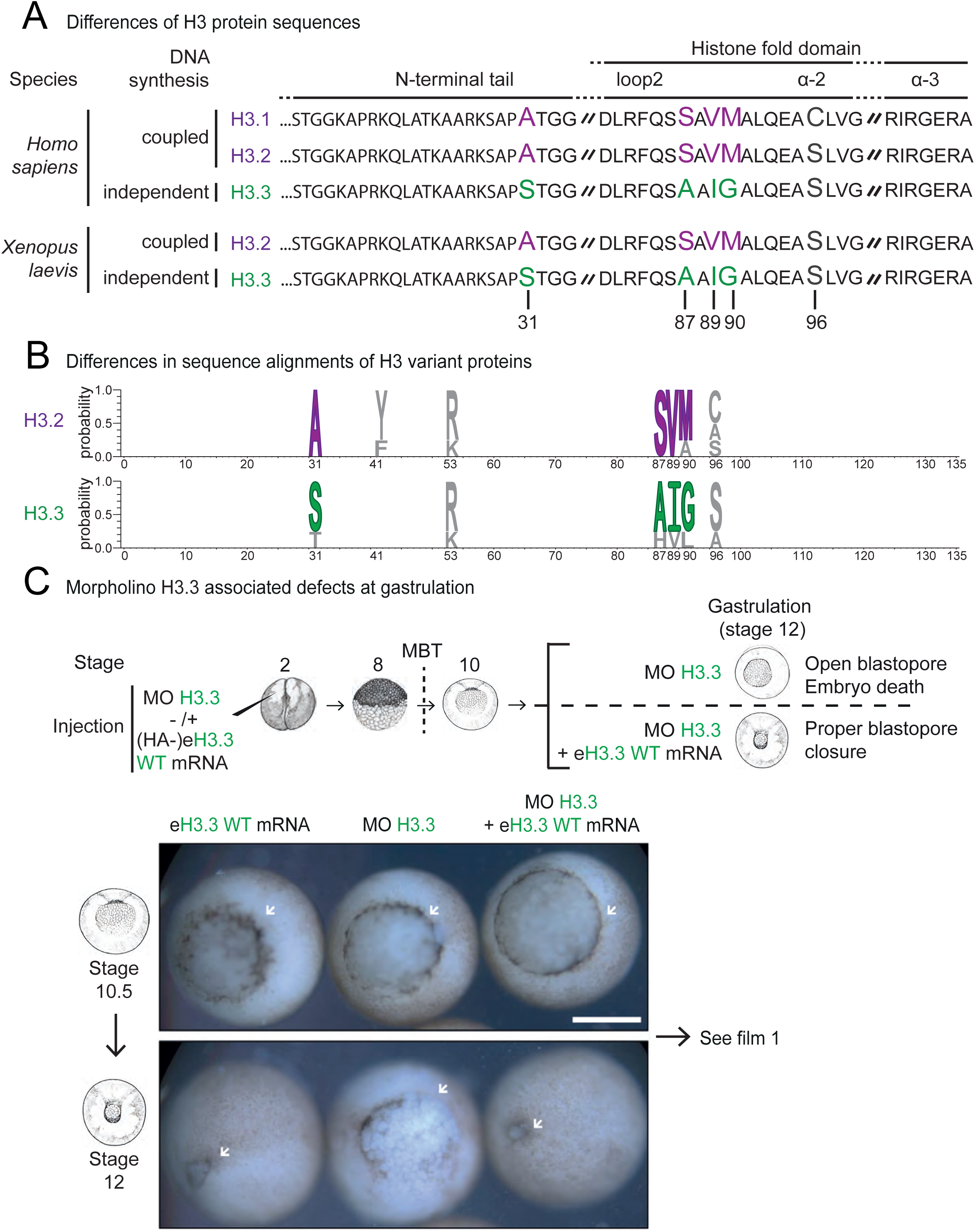
H3.3 is essential for gastrulation of *Xenopus laevis*. **A)** Best-studied non-centromeric H3 histone variants in *Homo sapiens* and *Xenopus laevis*. The two well-characterized forms of non-centromeric H3 variants correspond to the replicative histones H3.1 and H3.2 and the non-replicative histone H3.3, depicted in purple and green respectively. In human, the replicative H3 form is comprised of H3.1 and H3.2 that differ by only one residue at position 96, a cysteine and a serine respectively. The non-replicative form H3.3 shares more than 96% of identity with replicative forms, with five and four residue differences with H3.1 and H3.2 respectively. Additionally, *Xenopus laevis* embryos possess only one replicative histone variant, H3.2. Finally, histone sequences are conserved between *Homo sapiens* and *Xenopus laevis*. **B)** Logo for H3 variant sequence differences of *Homo sapiens, Mus musculus, Drosophila melanogaster, Xenopus laevis*, and *Arabidopsis thaliana* after performing multiple sequence alignment using MUSCLE. We display this alignment using WebLogo3. Specific histone variants are highlighted in green and purple. Histone variants show an extreme conservation between species. **C)** Developmental assay to monitor H3.3 functions. Morpholino and eH3.3 mRNA are injected at the 2-cell stage and associated defects can be observed at the gastrulation stage if there is no rescue. Scale bar corresponds to 500µm. See film 1.

### Permuting individually each amino acid of the H3.3 AIG motif towards H3.2 still ensures H3.3 functions during *Xenopus laevis* early development

To investigated the importance of the deposition pathway, we first look at the H3.3 histone chaperone recognition motif (Figure 2 A). Incorporation into chromatin of the non-replicative variant H3.3 occurs at any time of the cell cycle (DSI), and involves the HIRA complex in gene-rich regions (Ahmad and Henikoff 2002b; Ray-Gallet et al. 2002; Tagami et al. 2004; Ray-Gallet et al. 2011) and the DAXX/ATRX complex in heterochromatin regions (Gibbons et al. 1995; Yang et al. 1997; Drane et al. 2010; Goldberg et al. 2010; Lewis et al. 2010). In contrast, incorporation of the replicative variants is coupled to DNA synthesis (DSC) and is mediated by the CAF-1 complex (Stillman 1986; Smith and Stillman 1989; Gaillard et al. 1996; Tagami et al. 2004; Polo et al. 2006; Groth et al. 2007; Latreille et al. 2014). Importantly, structural studies enabled the identification of how histone chaperones discriminate the distinct histone variants through a motif located in the globular domain of histones (Elsasser et al. 2012; Liu et al. 2012; Ricketts et al. 2015). Both DAXX and HIRA complexes bind to the AIG motif of H3.3, with a particular affinity for the glycine at position 90. Therefore, we mutated H3.3 in the histone chaperone recognition motif to test their ability to rescue the loss of H3.3. We first designed single mutants for each residue of the AIG motif (Figure 2 B). To our surprise, every single mutant for the motif, *i*.*e*. eH3.3 A87S, eH3.3 I89V and eH3.3 G90M could rescue loss of H3.3, and embryo development occurred with the same efficiency. We verified that all constructs expressed at similar levels in the embryo (Supplemental figure 2 A) and incorporated into chromatin (Supplemental figure 2 B). Since it was possible that early development may allow looser interactions with the dedicated chaperones, we decided to completely change the recognition motif and assess both development and incorporation means.

**Figure 2:**
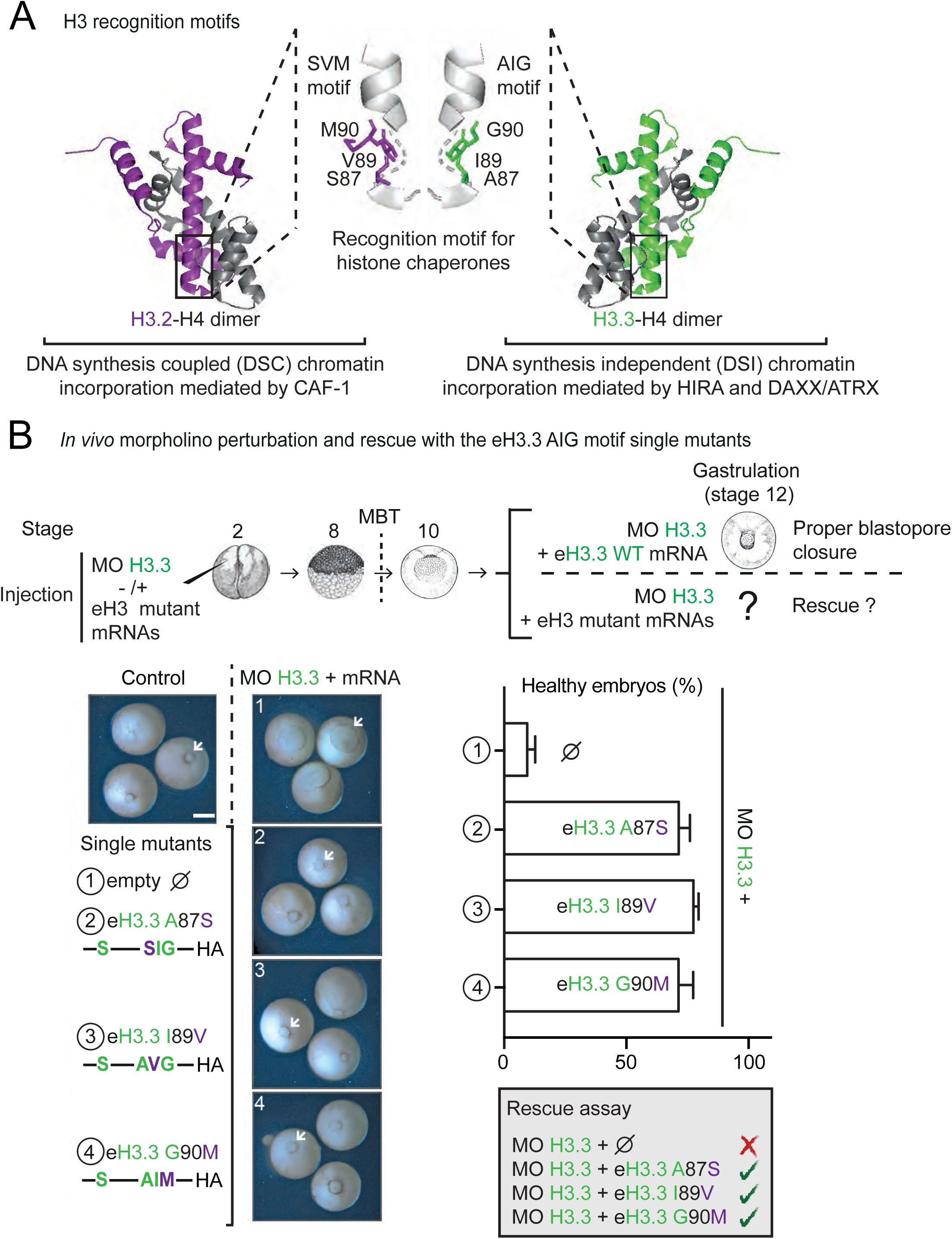
eH3.3 AIG single mutants rescue depletion of H3.3 during *Xenopus laevis* early development. **A)** Highlights of the histone chaperone recognition motif residues of H3 variants. Dedicated histone chaperones recognize histone variants by the H3.2 SVM and H3.3 AIG motifs. Although both motifs are structurally similar, the main difference appears for the residue 90 that is critical for histone chaperone binding. Crystal structure adapted from PDB ID codes: 5B0Z (Suzuki et al. 2016) and 3AV2 (Tachiwana et al. 2011). **B)** Rescue assays with H3.3 AIG single mutants. Injections are performed at 2-cell stage. The morpholino against H3.3 induces gastrulation defects that are rescued by all eH3.3 with single mutations of the AIG motif. Scale bar corresponds to 500µm. Quantification of properly developed embryos after injections of the different H3.3 AIG mutant forms shows no difference of rescue efficiency after H3.3 depletion. Each experiment has been reproduced at least 3 times with a minimum of 30 embryos.

### Swapping the H3.3 histone chaperone recognition motif with the one from H3.2 still enables *Xenopus laevis* early development yet switching the mode of incorporation

Since the individual substitution in the H3.3 motif did not affect the developmental rescue *in vivo*, we assessed whether mutation of all three residues of the motif would then affect H3.3 functions (Figure 3). To our surprise, in this context, this eH3.3 SVM hybrid form proved still able to rescue the loss of H3.3 as eH3.3 WT (wild-type), while eH3.2 WT cannot. In addition, such mutant form is expressed as well as incorporated into chromatin at similar levels in the embryo (Supplemental figure 3 A and B). Thus, by substituting the H3.3 recognition motif for its histone chaperone with the one from H3.2, we could ensure the rescue *in vivo*. Thus, the next question was whether in this context, the histone chaperone recognition motif actually determined the respective deposition pathways for each histone variant. The H3.3 dedicated chaperone complex HIRA recognized best H3 carrying the H3.3 AIG motif (Figure 4 A). However, p60, one subunit of the CAF-1 complex dedicated to H3.2, could perfectly recognize eH3.3 SVM, *i*.*e*. when the whole H3.3 motif is substituted to the one of H3.2. This shows that eH3.3 SVM can be recognized by CAF-1, arguing for a possible swap in the means for incorporation. We confirmed this finding *in vivo* by immunoprecipitation of eH3 forms directly from embryos at gastrulation stage (Supplemental figure 4 A). We found that p60 again only interacted with eH3 carrying the H3.2 SVM motif. Therefore, changing the motif alters the chaperone interactions. We thus explored the potential impact on the mode of histone variant incorporation. To test this aim, we performed chromatin assembly assays using extracts derived from *Xenopus laevis* eggs (Smythe and Newport 1991; Almouzni and Wolffe 1993). We supplemented Xenopus egg extracts with eH3.3 WT, eH3.2 WT or eH3.3 SVM and monitored their incorporation into chromatin using sperm nuclei under conditions allowing or not DNA synthesis (Figure 4 B). In interphase extracts, sperm DNA forms nuclei, can replicate and reassemble chromatin whereas mitotic extracts lack this DNA replication capacity. We isolated and analyzed sperm chromatin nuclei from interphase extracts, in presence or absence of the DNA synthesis inhibitor aphidicolin. Regardless of DNA synthesis, the incorporation of eH3.3 WT using sperm nuclei occurred with a similar efficiency in the presence or absence of DNA synthesis. By contrast, eH3.2 WT incorporation was severely affected by the presence of aphidicolin. Importantly, eH3.3 SVM incorporation shows the same dependency on DNA replication, arguing that its incorporation mode is switched toward a DNA-synthesis coupled mechanism. Consistently, p60 recruitment to chromatin is, as expected, highly reduced when DNA synthesis is inhibited. We further confirmed in mitotic extracts that only variant with the AIG motif could get incorporated independently of DNA synthesis (Supplemental figure 4 B). Based on these data, we conclude that the H3.3 histone variant could be efficiently provided regardless of the pathway for incorporation. Thus, defects at the time of gastrulation do not arise from a need for a distinct incorporation pathway outside S phase independently of DNA synthesis. It rather reflects an inherent feature of the variant when incorporated into chromatin. We thus examined more closely if the only remaining specific residue of H3.3, aminoacid 31, could account for this unique feature.

**Figure 3:**
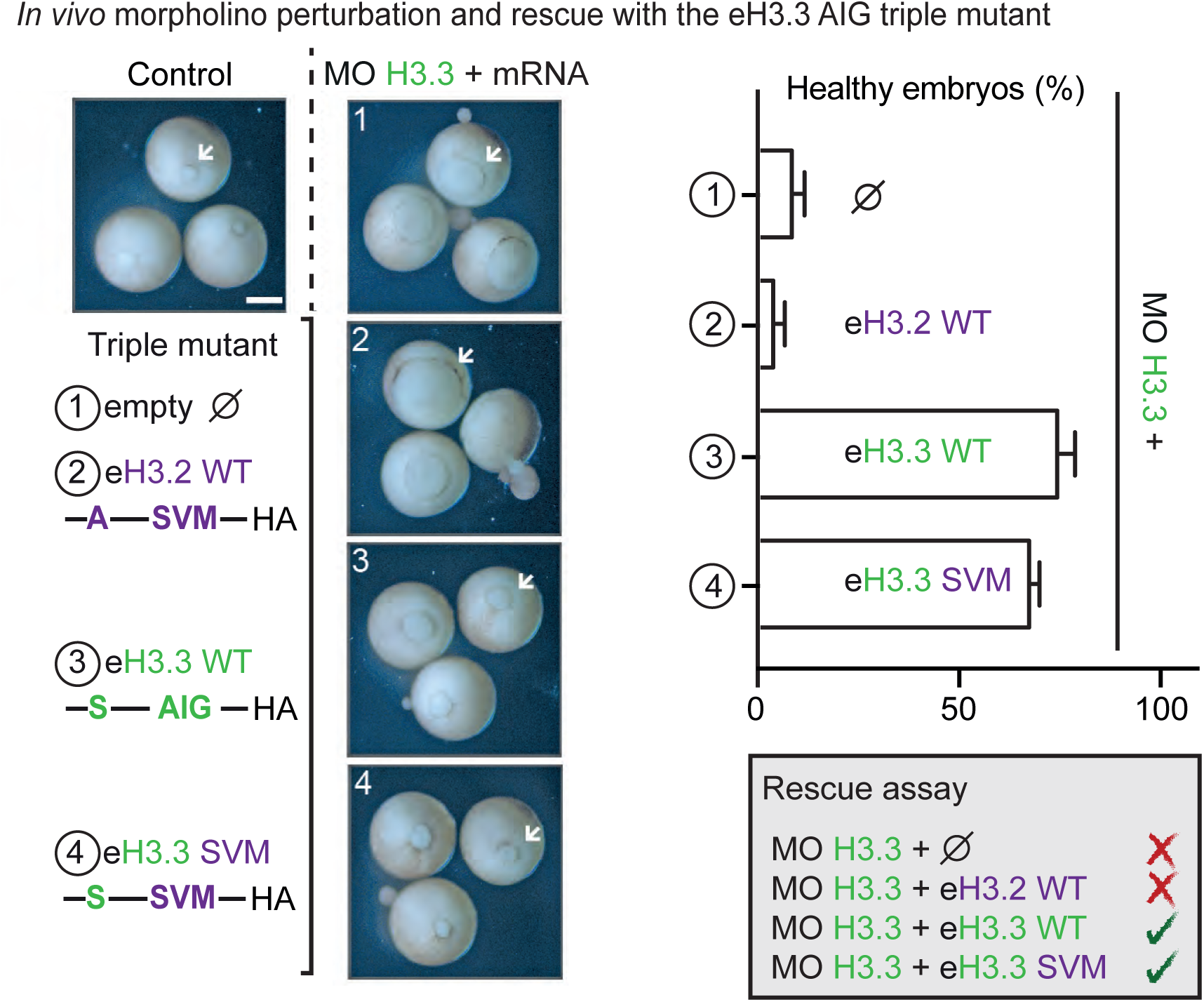
eH3.3 AIG triple mutant rescues depletion of H3.3 during *Xenopus laevis* early development. Rescue assays with eH3.3 triple AIG mutant. Injections are performed at 2-cell stage. The morpholino against H3.3 induces gastrulation defects that are not rescued by eH3.2. In contrast, a triple mutant of H3.3 for the AIG motif, eH3.3 SVM, rescues the phenotype while carrying the same SVM motif than H3.2. Scale bar corresponds to 500µm. Quantification of properly developed embryos after injection of the eH3.3 triple AIG mutant form shows similar rescue efficiency than eH3.3 WT after H3.3 depletion. Each experiment has been reproduced at least 3 times with a minimum of 30 embryos.

**Figure 4:**
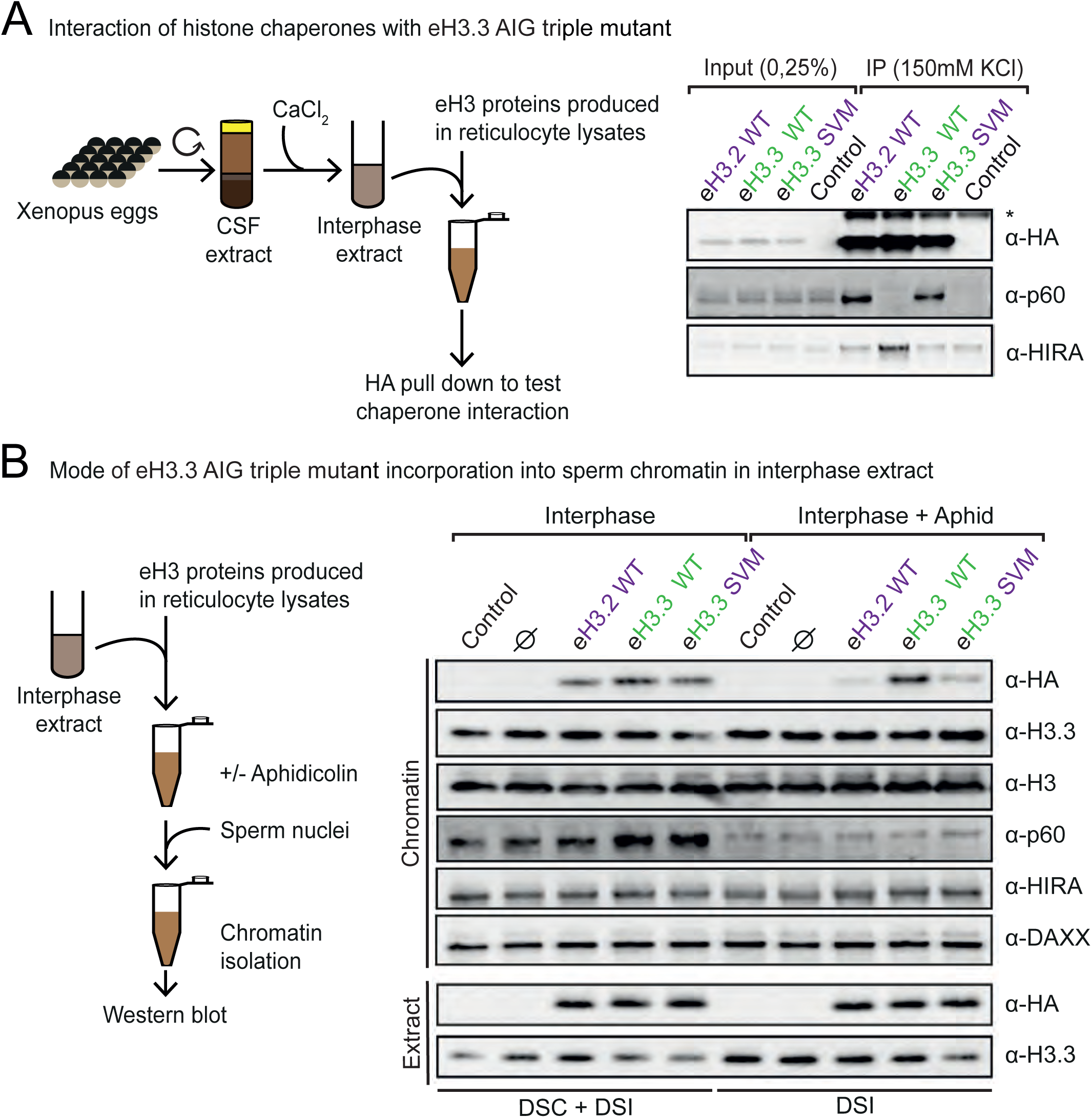
Swapping the AIG motif to the SVM motif leads to changes in chaperone interactions and histone modes of incorporation into chromatin *in vivo*. **A)** Immunoprecipitation of eH3 variant forms in interphase extract. Recombinant proteins are produced in rabbit reticulocyte lysates and pulled down by their HA-tag after incubation. HIRA only recognizes the AIG motif while p60 recognizes solely the SVM motif. * corresponds to a non-specific band. **B)** Incorporation of eH3 variant forms into sperm chromatin in interphase extract. Purified nuclei remodeled in the interphase extracts supplemented with indicated eH3.3 in the presence or absence of aphidicolin were analyzed by WB with indicated antibodies. While all eH3 variant forms are incorporated when both DSC and DSI pathways are available, only the eH3 variant forms with the AIG motif can be when only the DSI is available.

### H3.3S31 residue is critical for *Xenopus laevis* early development and is phosphorylated during the cell cycle

To address the role of the H3.3 residue at position 31, we first constructed a new H3.3 mutant, H3.3 S31A, containing an alanine instead of a serine at position 31 while keeping its original AIG motif (Figure 5 A and Supplemental figure 5 A). Remarkably, eH3.3 S31A was not able to rescue H3.3 depletion during Xenopus early development, indicating that the specific serine of eH3.3 at position 31 cannot be substituted by an alanine, which is notably the corresponding residue of H3.2, to fulfill H3.3 dedicated functions at this window of time during development. Interestingly, this particular residue has first been found phosphorylated in human cell line during mitosis (Hake et al. 2005). Furthermore, a threonine replaces this serine in Arabidopsis, a residue that can possibly also undergo phosphorylation, albeit this has not yet been documented. A key question was thus whether the actual need for H3.3 was linked to the capacity of H3.3S31 to be phosphorylated. To this end, we examined H3.3S31 phosphorylation in the Xenopus system using a Xenopus A6 cell line derived from kidney. By immunofluorescence, we detected a strong enrichment of H3.3S31p during mitosis (Figure 5 B). Interestingly, when compared to a mitotic modification, H3S10p, common to both H3 variant forms, its pattern was different, indicating a possible distinct function. While H3S10p covers the edges of mitotic chromosomes, H3.3S31p is enriched at centric and pericentric heterochromatin. This pattern, similar to previous observations in HeLa B cells (Supplemental figure 5 B) as in (Hake et al. 2005), is consistent with observations in mouse and monkey cell lines (Wong et al. 2009; Hinchcliffe et al. 2016). Kinases responsible for H3.3S31 phosphorylation are still unclear, currently implicating either CHK1 (Chang et al. 2015) or Aurora B (Li et al. 2017). Using Xenopus sperm chromatin, we further characterized the acquisition of this mark prior to or after replication in interphase extract or in extracts pushed into mitosis (Figure 5 C). We did not detect any significant signal for H3.3S31p in the soluble pool of H3.3 in either mitotic or interphase extracts, indicating that the modification is likely acquired once the variant is incorporated into chromatin. In addition, we could not detect any signal for this mark on non-remodeled sperm chromatin or on chromatin assembled in interphase extracts. This mark was most enriched in the fraction corresponding to isolated mitotic chromatin, in a profile similar to the H3S10p mark. We therefore conclude that H3.3S31p showed predominantly a mitotic chromatin-dependent mark in both somatic cells and reconstituted chromatin in early embryonic extracts, a mark imposed within chromatin and not prior to deposition. The next question was whether this modification or charge was also critical at the time of gastrulation in our rescue experiments.

**Figure 5:**
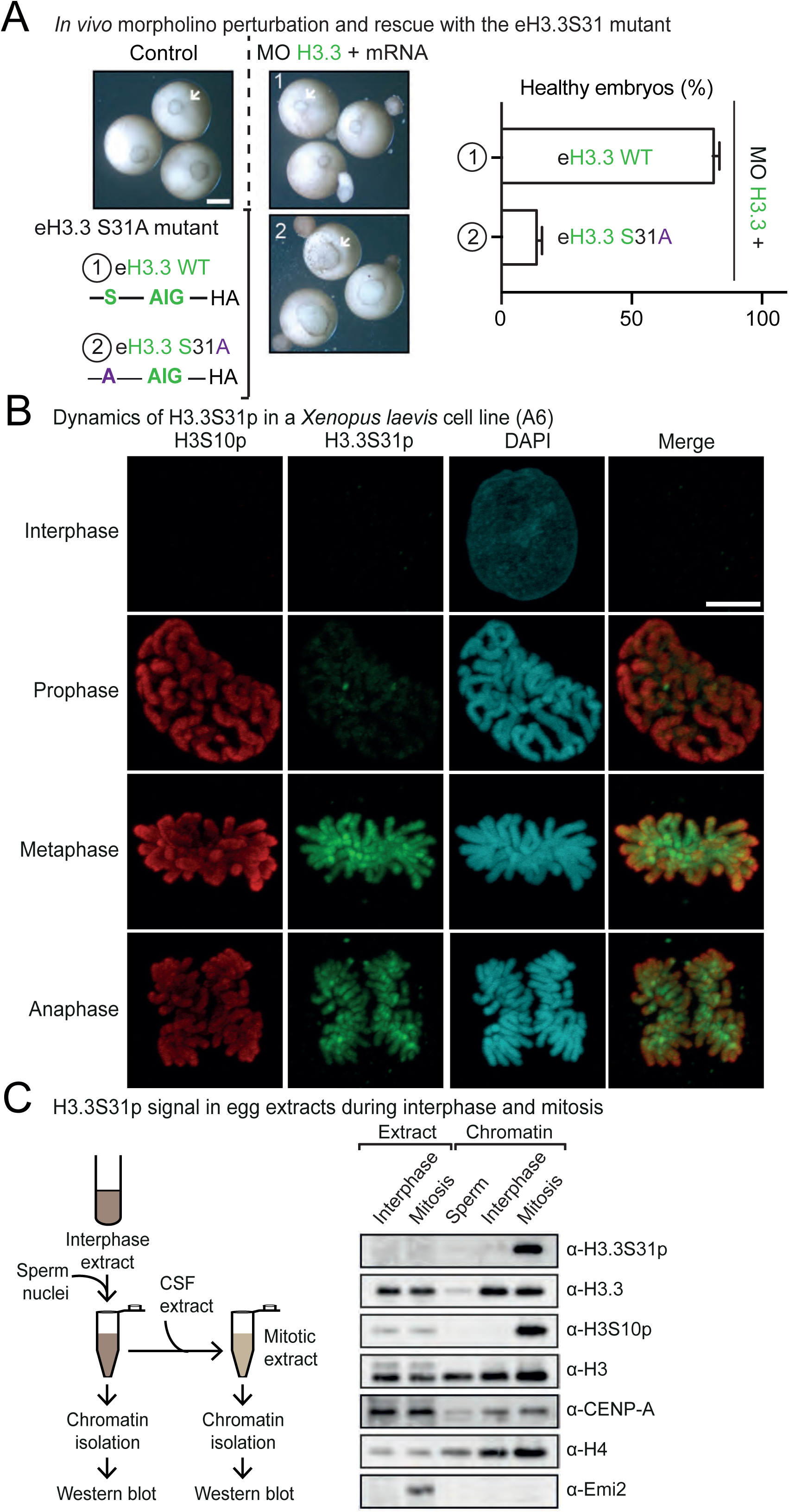
H3.3S31 is critical *Xenopus laevis* early development and is phosphorylated in this model organism. **A)** Rescue assays with eH3.3 S31A mutant. Injections are performed at 2-cell stage. Compared with eH3.3 WT, eH3.3 S31A mutant form cannot rescue the phenotype. Scale bar corresponds to 500µm. Quantification of properly developed embryos after injection of the eH3.3 triple AIG mutant form shows similar rescue efficiency than eH3.3 WT after H3.3 depletion. Each experiment has been reproduced at least 3 times with a minimum of 30 embryos. **B)** 3D-distribution and timing of H3.3S31 phosphorylation in A6 cell line. H3.3S31p follows the same dynamics than H3S10p, but does not localize to the same physical places. Scale bar represents 10 µm. **C)** Characterization of H3.3S31 phosphorylation in Xenopus cell-free extract. Non-remodeled sperm nuclei and nuclei purified after remodeling in interphase or mitotic extracts, H3 PTMs are analyzed by WB. H3.3S31p and H3S10p are enriched exclusively in sperm nuclei purified from mitotic extracts. The marks are prevalently found in the chromatin and not in H3.3 soluble form. Emi2 (Early mitotic inhibitor) serves as a control to confirm that extracts are in mitosis. CSF (Cytostatic factor).

### The negative charge of H3.3S31 is essential during *Xenopus laevis* early development at gastrulation

In order to explore a potential need for the phosphorylation of H3.3S31, we designed another H3.3 mutant for the serine carrying an aspartic acid, eH3.3 S31D, which acts as a phosphomimic version of this residue and carries a constitutive negative charge at position 31 that cannot be dynamically regulated by kinases and phosphatases (Figure 6 A). Remarkably, this mutant form could rescue the depletion of H3.3 comparable to the eH3.3 WT. Notably, all mutants were expressed and incorporated into chromatin in the same proportion (Supplemental Figure 6 A and B) and we verified that these mutations on the H3.3 tail did not indirectly alter the ability to interact with specific histone chaperones (Figure 6 B). Especially, neither H3.3 S31A nor S31D mutations affected the mode of incorporation of H3.3 in the Xenopus egg extracts-base chromatin assembly assays (Figure 6 C and Supplemental figure 6 C). This enabled us to discard any defects that could be related to inefficient incorporation. We can thus conclude that the residue at position 31 in H3.3, either as a serine or as a negatively charged residue (phosphomimic), is specifically needed for the function of the H3.3 variant once incorporated into chromatin as revealed at the time of gastrulation during early development in the vertebrate Xenopus.

**Figure 6:**
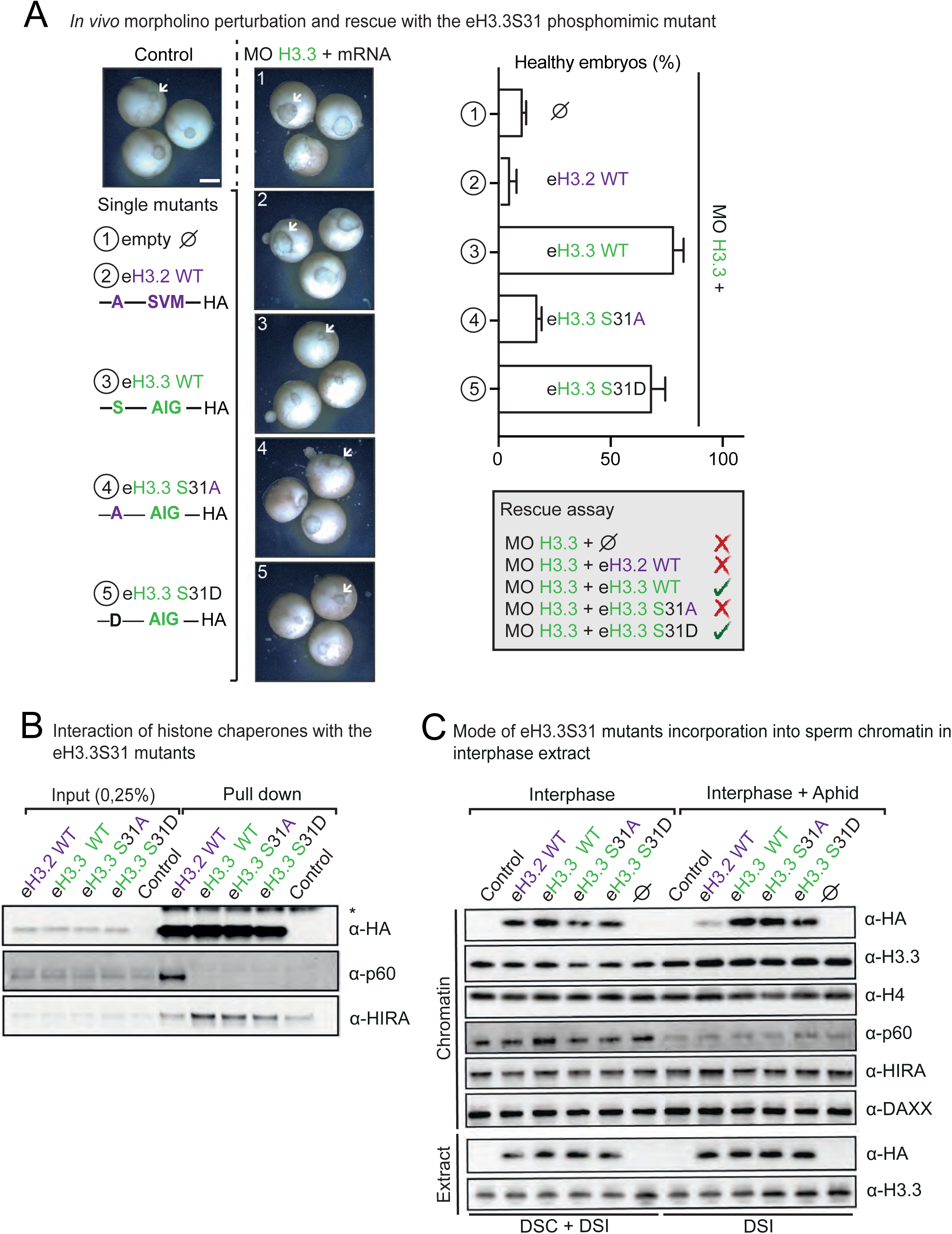
H3.3S31 negative charge is essential to rescue depletion of H3.3 during *Xenopus laevis* early development.

**A)** Rescue assays with eH3.3 S31D mutant. Injections are performed at 2-cell stage. In contrast to eH3.3 S31A, the use of an exogenous phosphomimic mutant, eH3.3 S31D, rescues, as the eH3.3 WT, the defects during gastrulation. Scale bar corresponds to 500µm. Quantification of properly developed embryos after injection of the H3.3 S31D mutant form shows similar rescue efficiency than eH3.3 WT after H3.3 depletion. In contrast, eH3.3 S31A mutant form leads to the same proportion of defects than eH3.2 after H3.3 depletion. Each experiment has been reproduced at least 3 times with a minimum of 30 embryos. **B)** Immunoprecipitation of eH3 S31 mutant forms in interphase extract. Recombinant proteins are produced in rabbit reticulocyte lysates and pulled down by their HA-tag after incubation. Mutations of the residue 31 do not alter histone chaperone interactions. * corresponds to a non-specific band. **C)** Incorporation of eH3 S31 mutant forms into sperm chromatin in interphase extract. Purified nuclei remodeled in the interphase extracts supplemented with indicated eH3.3 in the presence or absence of aphidicolin were analyzed by WB with indicated antibodies. Mutations of the residue 31 do not change the mode of incorporation of these forms.

## Discussion

By exploiting depletion/complementation assays in *Xenopus laevis* and monitoring the capacity to undergo gastrulation, we disentangle the critical role of H3.3 within chromatin through its aminoacid 31 independently of its mode of incorporation.

### H3.3 incorporation into chromatin is important during Xenopus early development regardless of its mode of incorporation

Replicative and non-replicative H3 variants do show distinct genome wide distributions (Clement et al. 2018), with enrichment of H3.3 at enhancers, or proximal to telomeres and centromeres as observed in several models in somatic cells and embryonic cells (Filipescu et al. 2014). How these distinct patterns arise and evolve dynamically during development and are then maintained in given lineages remains to be established. Moreover, how the unique properties of H3.3 influence cell cycle related functions and/or cell fate programming is still an open question. We first examined whether the mutations of the residues assigned to a key role in the choice of the deposition pathway (Ahmad and Henikoff 2002a; Elsasser et al. 2012; Liu et al. 2012; Ricketts et al. 2015) affect the capacity to rescue the depletion of H3.3 (Figure 2 and Figure 3). Surprisingly, an eH3.3 form containing H3.2 recognition motif still complements H3.3 functions in our developmental assay (Figure 3) even though this swap between motifs does effectively alter the chaperone interactions and the incorporation pathway (Figure 4). These data underline the fact that neither the chaperone interaction nor the mode of incorporation of the variant is critical to enable H3.3 specific roles at this time of development. Therefore, in the context of Xenopus early development, in sharp contrast with what one would have anticipated, it is the presence *per se* of H3.3 into chromatin that proves most important *in vivo*, regardless of the means to get incorporated, (Figure 7). This discovery shed light on unique roles performed by the single amino acid S31 on H3.3, highlighting the idea that *every amino acid matters* (Maze et al. 2014)!

**Figure 7:**
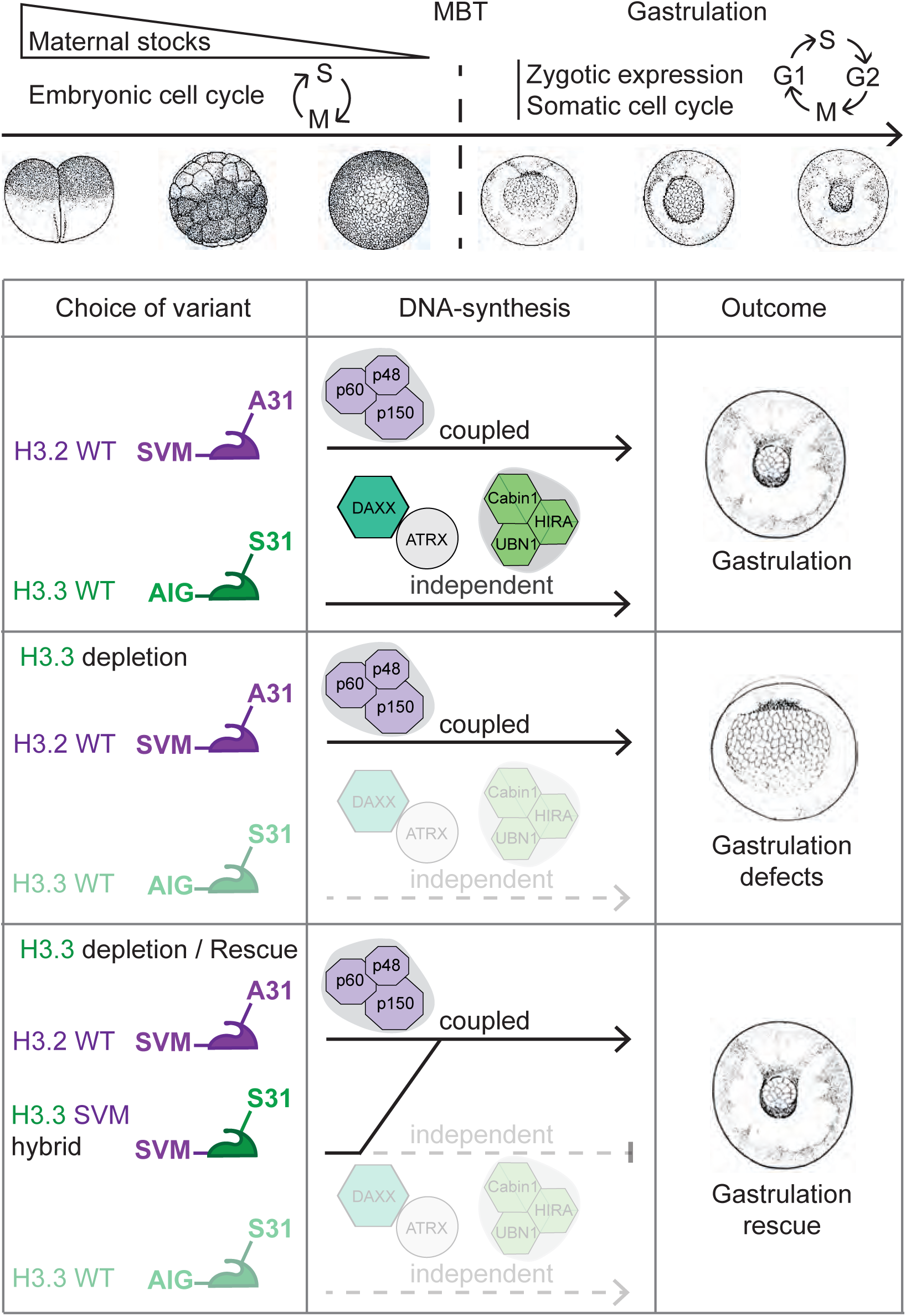
Graphical abstract. Defects associated with H3.3 depletion can be rescued with H3 histone variants carrying a potential negatively charge residue regardless of the mode of incorporation.

### H3.3S31 and a constitutive negative charge are essential for H3.3 functions during Xenopus early development

We showed by replacing the residue at position 31 in the N-terminal tail of H3.3 with the alanine found in H3.2 that such eH3.3 S31A mutant could not replace H3.3 (Figure 5 A). This proves that H3.3S31 is essential in our complementation assay after H3.3 depletion. Moreover, this serine on H3.3 can undergo phosphorylation while the alanine on H3.2 cannot. This specific phosphorylation of H3.3 regulated during the cell cycle shows enrichment during metaphase (Hake et al. 2005) (Figure 5 B). In our experiments, remarkably a phosphomimic form with a negative charge on the serine at position 31 could readily rescue the depletion of H3.3 (Figure 6 A). However, the exact role of H3.3S31 remains enigmatic. Although, we (Figure 5 and Supplemental figure 5) and others (Hake et al. 2005; Wong et al. 2009; Hinchcliffe et al. 2016) detected strong H3.3S31p signals primilary in mitosis, we cannot exclude that discrete sites of transcription could be marked but undetected in our assays. Indeed, this mark has been associated with particular transcribed regions in activated macrophages (Thorne et al. 2012). Furthermore, in mammals, a function for H3.3S31p has been linked to intron retention and pre-mRNA processing by preventing binding of the tumor suppressor BS69, also known as ZMYND11 (Zinc finger MYND domain-containing protein 11), an H3.3K36me3-specific reader (Guo et al. 2014; Wen et al. 2014). Our assays demonstrate the importance of H3.3S31 at gastrulation (Figure 5 and Figure 6), a critical time for lineage commitment accompanied by dramatic cell cycle changes and modifications in gene expression patterns. Taking into account the specific importance given to H3.3 that ranges from transcription to reprogramming (Ahmad and Henikoff 2002b; Schwartz and Ahmad 2005; Ng and Gurdon 2008; Jullien et al. 2012), the observed developmental defects could arise from a role for H3.3S31 and its phosphorylation in transcription initiation or maintenance. Importantly, MBT in Xenopus also leads to changes in the cell cycle progression: establishment of the longer cell cycle “somatic-type” with the gap phases and acquisition of checkpoints (Etkin 1988; Masui and Wang 1998). Moreover, histones are key component in regulating the start of the MBT, possibly through titration mechanisms (Almouzni and Wolffe 1995; Amodeo et al. 2015). It should be valuable to explore the role of specific variants into this context. Thus, H3.3S31 function could equally depend on a specific role of H3.3 role during the cell cycle. Indeed, H3.3 has been involved in chromosome segregation as shown in double H3.3 KO embryonic stem cells that leads to an increase in anaphase bridges and lagging chromosomes (Jang et al. 2015). Furthermore, this mark can coat lagging chromosomes in parallel to p53 activation in order to prevent aneuploidy (Hinchcliffe et al. 2016). Considering H3.3S31p accumulation at centromeres in mitosis (Figure 5 B) (Hake et al. 2005), it is thus tempting to speculate that H3.3S31p could crosstalk with CENP-A incorporation at late mitosis and the beginning of G1. Interestingly, CENP-A incorporation at the centromere has been shown to be dependent on both the presence of H3.3 as a placeholder (Dunleavy et al. 2011) and on transcription (Ohkuni and Kitagawa 2011; Bobkov et al. 2018). To reconcile both aspects, we could envision that H3.3S31p may act as a phosphoswitch in mitosis to regulate transcription at critical chromosomal landmarks and in interphase to control/maintain a transcription program. Alternatively, one could also consider that the alanine present on H3.2 could prevent particular interactions specific to H3.3S31 and permitted by an aspartic acid. Notably in Arabidopsis, the replicative form cannot be rescued by H3.3 (*i*.*e*. A31T mutation) (Jiang and Berger 2017). Indeed, ATXR5 (Arabidopsis trithorax-related protein 5), a plant specific H3K27 methyltransferase, specifically recognizes H3.1 and not H3.3 (Jacob et al. 2014). Therefore, the alanine at position 31 on replicative histone H3.1 is prevents the heterochromatinization of H3.3-rich regions during replication. Thus, in plants, the replicative histone H3 variant avoids the presence of a negative charge at position 31. In our work, we show that there is potentially no need for the dynamics of the mark. Rather, we demonstrate a need to provide a negative charge or to avoid an alanine during early development. Indeed, our data indicate that the dynamics of the H3.3S31p may be dispensable, since the phosphomimic mutant is able to fully rescue H3.3 depletion (Figure 6). We can foresee several possible explanations. On the one hand, the negative charge at position 31 could be an absolute requirement for H3.3 functions regardless of any dynamics. On the other hand, the negative charge on this residue could be associated with mechanisms that occur solely in mitosis and the dynamics may simply be ensured by removal of the variant without invoking a particular phosphatase. More work, specifically exploring eviction mechanisms is needed to disentangle these two possibilities. Thus, the residue at position 31 of the replicative and non-replicative H3 variant may be key to promote/allow or exclude the presence of a negative charge with different outcomes for critical binding partners. Conversely, other modifications on H3 tail may impact S31 and in this respect the existence of mutations affecting neighboring residues will be interesting to explore. Notably, H3.3S31 is close to residues often found substituted in aggressive cancers like pediatric glioblastoma (H3 K27M and G34R) (Heaphy et al. 2011; Jiao et al. 2011; Schwartzentruber et al. 2012; Wu et al. 2014). Thus, it will be very interesting to evaluate the connection between these particular cancers involving oncohistones with this H3.3 specific phosphorylation. Altogether, we show that H3.3S31 is the key residue that confers specific functions to H3.3 within chromatin. It establishes for the first time the importance of a distinct histone variant residue for the proper development of a vertebrate during gastrulation. Future work should explore whether a similar requirement also occurs in mammals to provide a comprehensive view of the importance of the non-replicative variant H3.3 and its role during vertebrate development and in disease states.

> “*Qu’importe le flacon, pourvu qu’on ait l’ivresse*”, Alfred de Musset

## Material & Methods

### *Xenopus laevis* embryo manipulation

We used *Xenopus laevis* adults (2 years old) from the Centre de Ressource Biologie Xenope. We prepared embryos at 18°C as in (Almouzni et al. 1994) and staged them according to (Nieuwkoop and Faber 1994). We acquired embryos during gastrulation with a MZFLIII magnifier (Leica) and the SPOT software. Animal care and use for this study were performed in accordance with the recommendations of the European Community (2010/63/UE) for the care and use of laboratory animals. Experimental procedures were specifically approved by the ethics committee of the Institut Curie CEEA-IC #118 (Authorization APAFIS#11226-2017091116031353-v1 given by National Authority) in compliance with the international guidelines. David Sitbon, Ekaterina Boyarchuk and Geneviève Almouzni possess the Authorization for vertebrates’ experimental use.

### Plasmid cloning and mRNA transcription

We cloned all H3 cDNAs in the pβRN3P vector (Zernicka-Goetz et al. 1996). This vector stabilizes RNA and improves their translation efficiency while injected into Xenopus eggs. In addition, an HA tag is inserted in the C-terminal of H3. We obtained mRNAs by *in vitro* transcription of PCR-amplified fragments of the different pβRN3P vectors (Forward: 5’-gtaaaacgacggccagt-3’ and Reverse: 5’-ggaaacagctatgaccatga-3’). We transcribed mRNAs starting with 5ng of PCR-amplified fragment, 10µL Buffer 5X, 5µL of DTT 100mM, 0.25µL bovine serum albumin (BSA) 10mg/mL (NEB), 5µL of ATP, CTP, UTP 10mM, 1.65µL GTP 10mM (Sigma-Aldrich), 3.35µL Me7GTP 10mM (NEB), 2µL of RNasin Plus RNase Inhibitor (Promega), and 50µL H2O qsp. After 10min on ice, we added 2µL of T3 RNA Polymerase (Promega) to each sample as well as after 30min at 37°C for another 1h30. After DNA digestion, we extracted mRNAs with phenol-chloroform and purified them through Sephadex G-50 Quick Spin Column for radiolabeled RNA purification (Sigma-Aldrich), previously equilibrated 6 times with 1mL of TE 10mM.

### Morpholino and mRNA microinjection into *Xenopus laevis* embryo

We microinjected 2-cell embryos using a Brinkmann micromanipulator and a Drumond microinjector on two-cell stage eggs with an injection volume set to 9.2nL to deliver the appropriate quantity of morpholino and mRNAs (1X=4.6ng). Between 20 and 30 embryos are injected per condition, with at least 3 biological replicates. We knocked down H3.3 with morpholino designed to bind to the initiation region of *Xenopus laevis* H3.3 mRNAs (Szenker et al. 2012).

### *Xenopus laevis* embryo protein extract preparations and Western blotting

We prepared total protein extracts from *Xenopus laevis* embryos using the CelLytic Express reagent (Sigma-Aldrich) and centrifuging at full speed the extracts after 30min incubation. We analyzed protein samples by electrophoresis on 4%–12% NuPAGE SDS-PAGE gels with MES SDS Running buffer (Life Technologies) and corresponding LDS buffer NuPage (Invitrogen) with DTT. Primary antibodies were detected using horse-radish-peroxidase-conjugated secondary antibodies (Jackson Immunoresearch Laboratories) and ECL2 kit (Pierce).

### *Xenopus laevis* embryo immunoprecipitation

We collected embryos at stage 12 and centrifuged them at 25000g after 3 washes in TE buffer (10mM Hepes, 75mM KCl and 50mM Sucrose). After ultracentrifugation at 150000g of the supernatant, we used 100µg of each condition for every immunoprecipitation overnight at 4°C in 400µL of IP buffer (10mM Hepes, 150mM KCl, 0;05% NP-40, 5mM DTT and protease inhibitors). After 3 washes, we eluted proteins and analyzed samples by electrophoresis on 4%–12% NuPAGE SDS-PAGE gels with MOPS SDS Running buffer (Life Technologies) and corresponding LDS buffer NuPage (Invitrogen) with DTT. Primary antibodies were detected using horse-radish-peroxidase-conjugated secondary antibodies (Jackson Immunoresearch Laboratories) and ECL2 kit (Pierce).

### *Xenopus laevis* embryo fractionation

We collected 40 embryos at stage 12 and performed fractionation as in (Szenker et al. 2012). We lysed the embryos in 200µL of Lysis Buffer 1 (10mM Tris-HCl pH 7.5, 200mM NaCl, 5mM MgCl2, 0.5% NP40, protease inhibitors) or Lysis Buffer 2 (10mM Tris-HCl pH 7.5, 10mM NaCl, 5mM MgCl2, 0;5% NP40, protease inhibitors) and centrifuged them at 1000g for 2min at 4°C. We obtained the clear soluble fraction after ultracentrifugation of the supernatant at 160000g for 1 hour at 4°C from Lysis Buffer 1. We washed the chromatin fraction from Lysis Buffer 2 after centrifugation in Buffer A (10mM Tris-HCl pH 7.5, 15mM NaCl, 60mM KCl, 0.34M sucrose, 1mM DTT, protease inhibitors). We performed MNase (Roche Diagnostics, #1010792100) digestion during 10min with 11.25U/mL final concentration, after addition of CaCl2 (2mM final) and RNase A (Roche Diagnostics, #10109142001) treatment (75µg/ml final, 5min at 37°C). We stopped the reaction by EDTA (50mM final) and recovered solubilized chromatin fraction after a centrifugation at 1000 g for 2min.

### *Xenopus laevis* egg extract preparation

We prepared *Xenopus laevis* sperm nuclei and low-speed extracts arrested by CSF (cytostatic factor) of *Xenopus laevis* eggs as previously described (Kornbluth et al. 2001). Briefly, we collected eggs freshly and centrifuged them at low speed (16000g) to conserve the mitotic phase, below lipids. We induced interphase by the addition of CaCl_2_ at the final concentration 0.06mM to CSF-arrested egg extracts. We added sperm chromatin at a concentration 1000– 4000 nuclei/µl. After DNA replication, a 2/3 volume of the corresponding CSF-arrested extract was added to induce mitosis.

### *Xenopus laevis* egg extract immunoprecipitation

Briefly, we produced recombinant H3 mutant proteins from mRNAs using rabbit reticulocyte lysate (Promega L4600). After 3h of incubation at 4°C in interphase extracts followed by another 3h incubation with anti-HA beads, we pulled-down and washed the proteins in 0.8x CSF-XB buffer (10mM Hepes-KOH, pH 7.7, 100mM KCl, 2mM MgCl2, 5mM EGTA) containing 5% glycerol, 0.5% Triton X-100 and protease and phosphatase inhibitors.

### *Xenopus laevis* sperm chromatin purification

We diluted fivefold 100µl aliquots of each reaction with 0.8× CSF-XB buffer containing 20mM β-glycerophosphate, 5% glycerol and 0.5% Triton X-100, which we incubated for 1 min at RT. We then layered the samples onto a 35% glycerol-containing CSF-XB cushion and centrifuged them at 10000g for 5 min at 4°C. We resuspended the pellets in the same buffer, and repeated the centrifugation step. For purification of interphase chromatin, we diluted 100µl aliquots of extract with 0.8× CSF-XB buffer containing 20 mM β-glycerophosphate and 5% glycerol, which we incubated for 1 min at RT, followed by centrifugation through the cushion at 10000g for 5 min at 4°C. We analyzed protein samples by electrophoresis on 4%–12% NuPAGE SDS-PAGE gels with MES SDS Running buffer (Life Technologies) and corresponding LDS buffer NuPage (Invitrogen) NuPage reducing agent (Invitrogen). Primary antibodies were detected using horse-radish-peroxidase-conjugated secondary antibodies (Jackson Immunoresearch Laboratories). We revealed the signal by chemiluminescence with SuperSignal West Dura Extended Duration Substrate (ThermoFisher). The signal was acquired using ChemiDoc Imager (Biorad).

### Chromatin assembly assays

We added 100000 sperm nuclei to each 150µL of corresponding extracts with or without aphidicolin (50µg/mL). We then supplied 15µL of the rabbit reticulocyte lysate used to produce recombinant H3 mutant proteins from mRNAs. After 40min of incubation at room temperature, we purified chromatin and we analyzed protein samples by electrophoresis as described above.

### WebLogo sequence alignment

After performing multiple sequence alignment using MUSCLE (Edgar 2004), we displayed the alignments using WebLogo3 (Crooks et al. 2004) with probability units.

### Antibody list

| Antibody List | WB | IF |
| --- | --- | --- |
| HA (3F10, Roche) | 1/1000 | / |
| H3.3S10p (Mabi0312, Wako) | 1/1000 | 1/1000 |
| H3.3S31p (ab92628, Abcam) | 1/1000 | 1/200 |
| H3.3 (M01, Abnova) | 1/200 | / |
| H4 (05-858, Upstate) | 1/1000 | / |
| Tubulin (T9026, Sigma) | 1/1000 | / |
| H3 (ab1791, Abcam) | 1/5000 | / |
| xEmi2 (From Mary Dasso) | 1/1000 | / |
| xHIRA (Agro Bio) | 1/1000 | / |
| xDAXX (From Mary Dasso) | 1/500 | / |
| xp60 (Agro Bio) | 1/1000 | / |
| xCENP-A (From Aaron Straight) | 1/1000 | / |

### Immunofluorescence and epifluorescence microscopy

We first fixed A6 and HeLa B cells on coverslips for 20 min in 4% and 2% paraformaldehyde, respectively, before to permeabilize them with 0.2% Triton X-100. We blocked them for 45min with 5% bovine serum albumin. We then incubated coverslips with primary and secondary antibodies and stained them with DAPI. We mounted the coverslips in Vectashield medium. We used a Confocal Zeiss LSM780, and we acquired images using 63×/1.4NA under Zen blue software (Zeiss Germany).

## Supporting information

Film 1

Film 2

## Acknowledgments

We are grateful to Dominique Ray-Gallet, Shauna Katz and Jean-Pierre Quivy for critical reading and to members of UMR3664 for helpful discussions. We thank Mary Dasso and Aaron Straight for Xenopus histone chaperone antibodies. This work was supported by la Ligue Nationale contre le Cancer (Equipe labellisée Ligue), ANR-11-LABX-0044_DEEP and ANR-10-IDEX-0001-02 PSL, ANR-12-BSV5-0022-02 “CHAPINHIB”, ANR-14-CE16-0009 “Epicure”, ANR-14-CE10-0013 “CELLECTCHIP”, EU project 678563 “EPOCH28”, ERC-2015-ADG-694694 “ChromADICT”, ANR-16-CE15-0018 “CHRODYT”, ANR-16-CE12-0024 “CHIFT”, “Parisian Alliance of Cancer Research Institutes”. DS was supported by PSL.

## Author contributions

D.S., E.B. and G.A. conceived the overall strategy and wrote the paper. G.A. supervised the work. D.S. performed *Xenopus laevis* and cellular experiments and analyzed data. E.B. performed *Xenopus laevis egg* extract experiments and analyzed data. Critical reading and discussion of data involved all authors.

---

**Film 1: Endogenous H3.3 depletion leads to gastrulation defects and stops *Xenopus laevis* early development.** Left embryo is injected with eH3.3 WT mRNA while the middle embryo is injected with 4.6ng of morpholino against H3.3. Right embryo is injected with both morpholino and eH3.3 WT mRNA. Injection of the morpholino leads to a gastrulation defects followed by developmental arrest that can be rescued by co-injection of eH3.3 WT mRNA.

**Film 2: Dose-dependent H3.3 depletion defects.** From left to right, the three embryos have been injected with 4.6ng, 9.2ng, 18.4ng of morpholino against H3.3. Right embryo has not been injected. Two replicates for each condition are showed. Injection of increased morpholino concentration leads to earlier gastrulation defects than previously observed.

**Supplemental figure 1:**
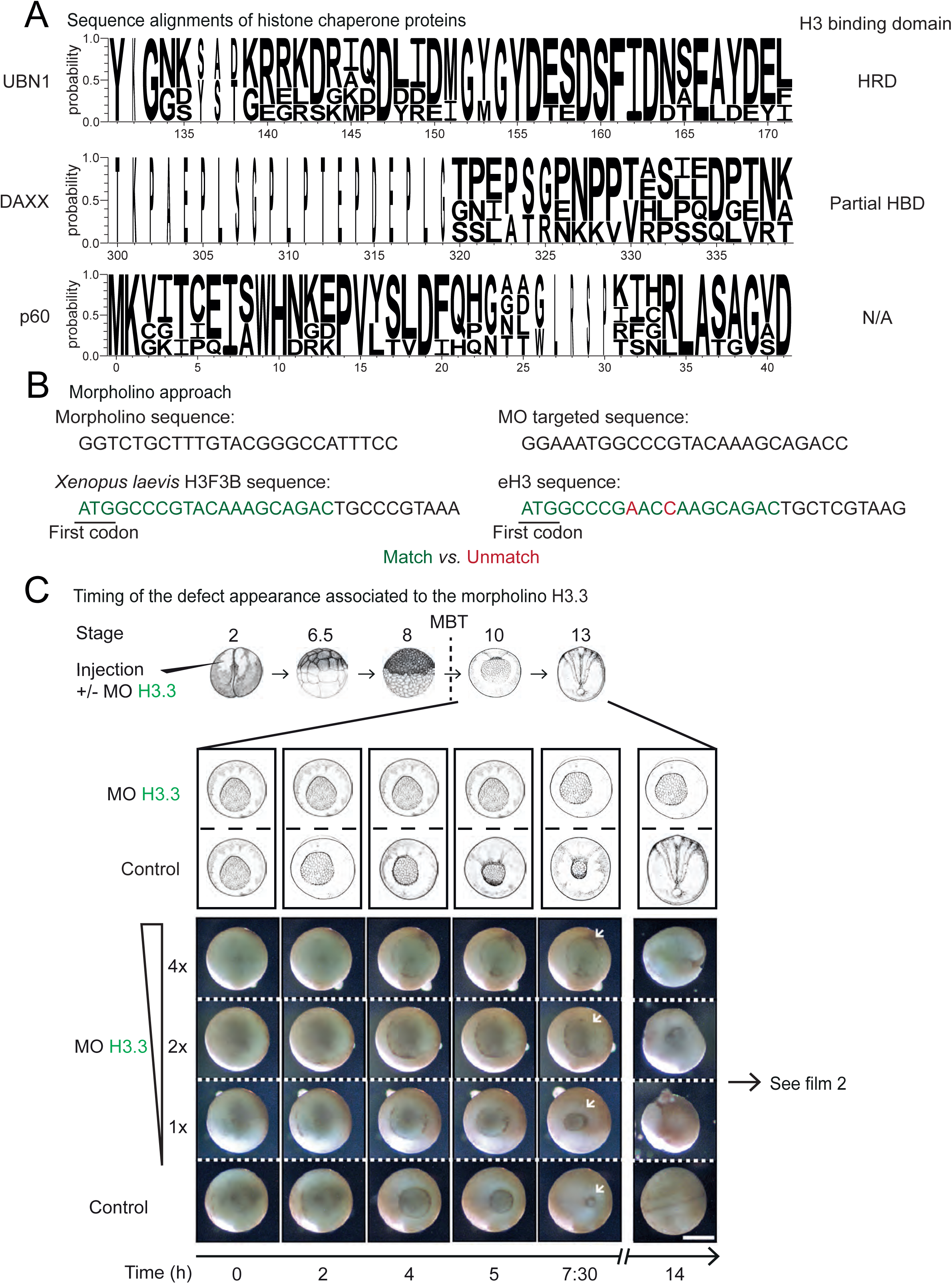
Histone chaperone sequence conservation and morpholino strategy and titration. **A)** Logo for UBN1, DAXX and p60 partial sequences of *Homo sapiens, Mus musculus, Drosophila melanogaster, Xenopus laevis* after performing multiple sequence alignment with MUSCLE. Complete histone-binding domain HRD (131-171) of UBN1 is presented, while only a part (from 300 to 339) of the DAXX histone-binding domain HBD (178-389) is displayed. No crystallographic data are available regarding the CAF1 complex but the overall sequence shows variation as exemplified by the 41 first residues. Histone chaperones show little conservation between species. Final alignment was made using WebLogo3. **B)** Sequences of the morpholino against endogenous H3.3 and its targeted sequence. The corresponding sequences matching the targeted sequence are highlighted in green for the first 10 codons of *Xenopus laevis* H3F3B and eH3 sequences. In contrast, differences with the targeted sequence are emphasized in red. **C)** Titration of the morpholino against H3.3. Increased concentrations of morpholino lead to earlier developmental defects. Morpholino against H3.3 is injected at the 2-cell stage with 3 different doses. While the lower one previously used in (Szenker et al. 2012) induces a defect at late gastrulation, two and four times more morpholino induce a defect appearing during early gastrulation. See film 2.

**Supplemental figure 2:**
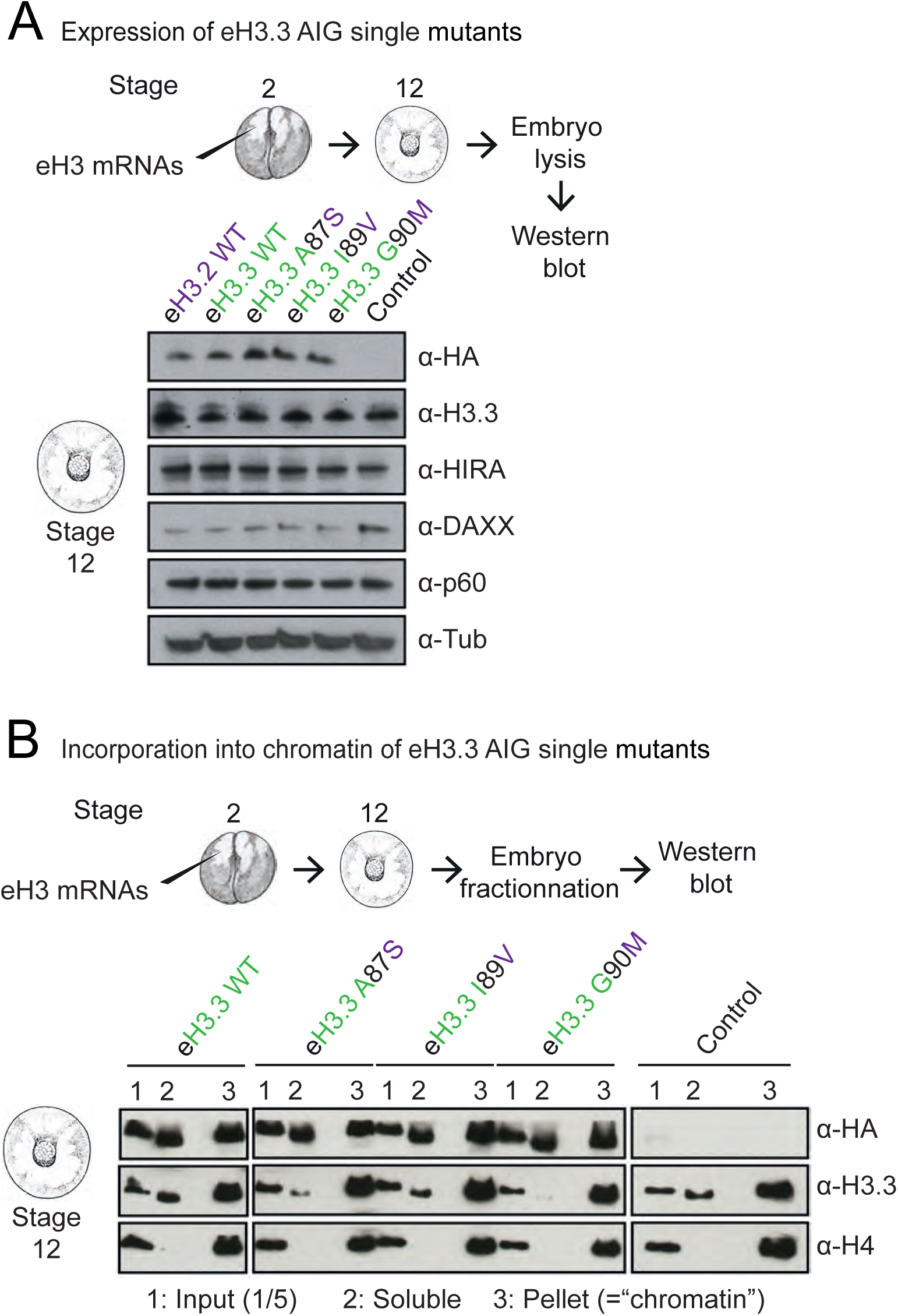
Single mutant forms of eH3.3 AIG motif are all expressed and incorporated in a similar fashion *in vivo*. **A)** Western blot to control eH3.3 AIG single mutant expression in embryos. Every eH3.3 AIG single mutants are expressed at similar levels in whole embryo extracts at stage 12. **B)** Western blots to control eH3.3 AIG single mutant incorporation in embryos. Fractionation of stage 12 embryos shows that all mutants are insoluble at 600nm NaCl, and therefore most likely incorporated into chromatin. Note that both eH3.3 WT and Control conditions are reused in Supplemental Figure 3 B and 6 B since each condition used comes from the same western blot.

**Supplemental figure 3:**
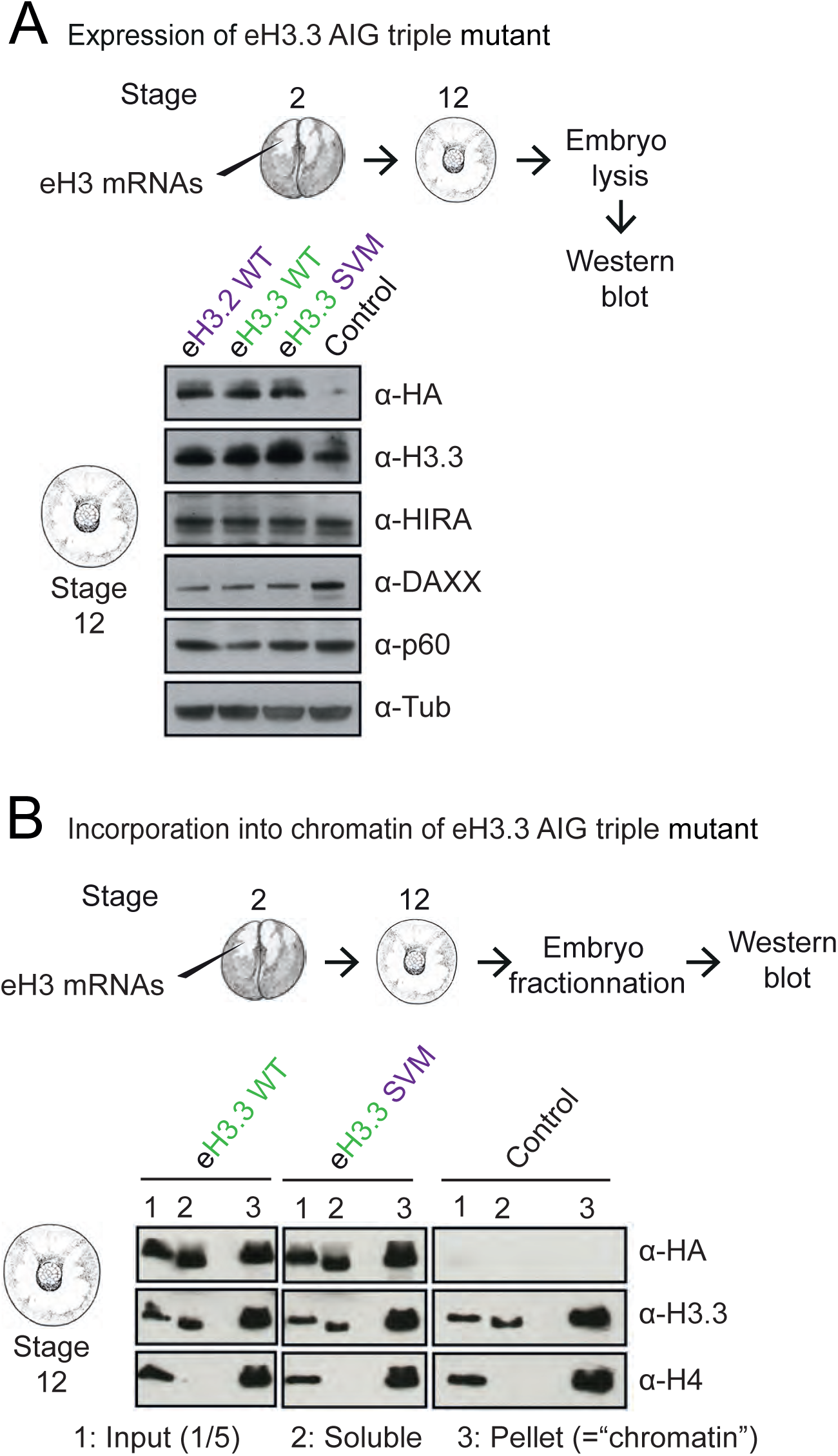
Triple mutant form of eH3.3 AIG motif is efficiently expressed and incorporated *in vivo*. **A)** Western blot to control eH3.3 AIG triple mutant expression in embryos. eH3.3 AIG triple mutant expression compares with the other eH3 conditions in whole embryo extracts at stage 12. **B)** Western blots to control eH3.3 AIG triple mutant incorporation in embryos. Fractionation of stage 12 embryos shows that this mutant remains insoluble at 600nm NaCl, and therefore most likely incorporated into chromatin. Note that both eH3.3 WT and Control conditions are reused in Supplemental Figure 2 B and 6 B since each condition used comes from the same western blot.

**Supplemental figure 4:**
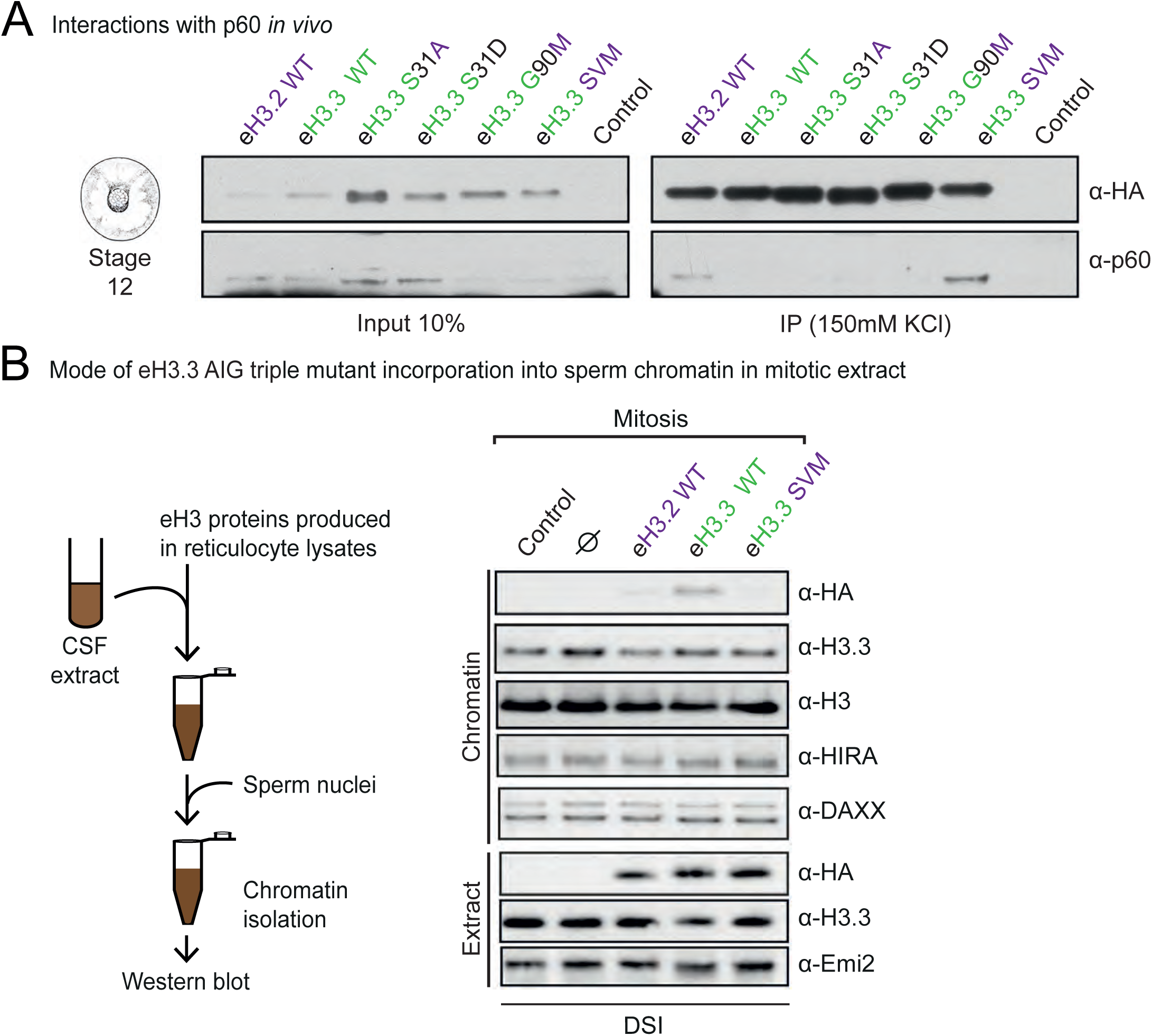
p60 histone chaperone interaction *in vivo* and mode of incorporation of eH3.3 AIG triple mutant in mitotic extract *in vitro*. **A)** Immunoprecipitation of eH3 variant forms from stage 12 embryos. Only the SVM motif is recognized by p60 *in vivo*. **B)** Incorporation of eH3 variant forms into sperm chromatin in mitotic extract. Purified nuclei remodeled in the CSF-arrested extracts supplemented with eH3.3 mutants were analyzed by WB with indicated antibodies. In mitotic extract where the only pathway available is the DSI, only the eH3 variant form with the AIG motif, *i*.*e*. eH3.3 WT, is incorporated.

**Supplemental figure 5:**
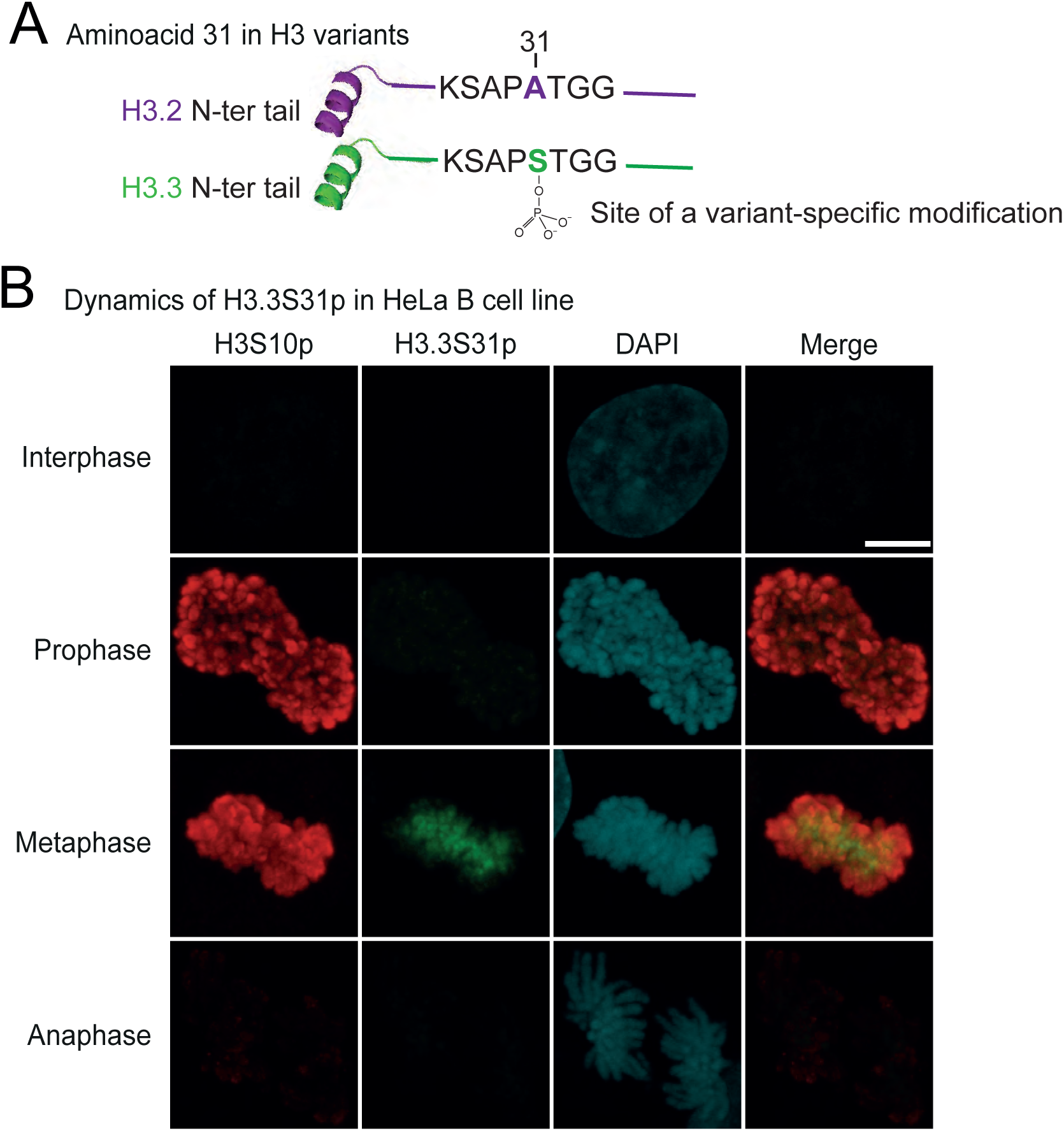
H3.3S31 phosphorylation in human has the same dynamics that in *Xenopus laevis*. **A)** Highlights of the histone tail specific residue of H3 variants. H3.3 tail possesses a serine at position 31 that can be phosphorylated, while the alanine of H3.2 cannot. Crystal structure adapted from PDB ID codes: 5B0Z (Suzuki et al. 2016) and 3AV2 (Tachiwana et al. 2011). **B)** 3D-distribution and timing of H3.3S31 phosphorylation in HeLa B cell line. As in the Xenopus, H3.3S31p follows the same dynamics than H3S10p, but does not localize to the same physical places. Scale bar represents 10 µm.

**Supplemental figure 6:**
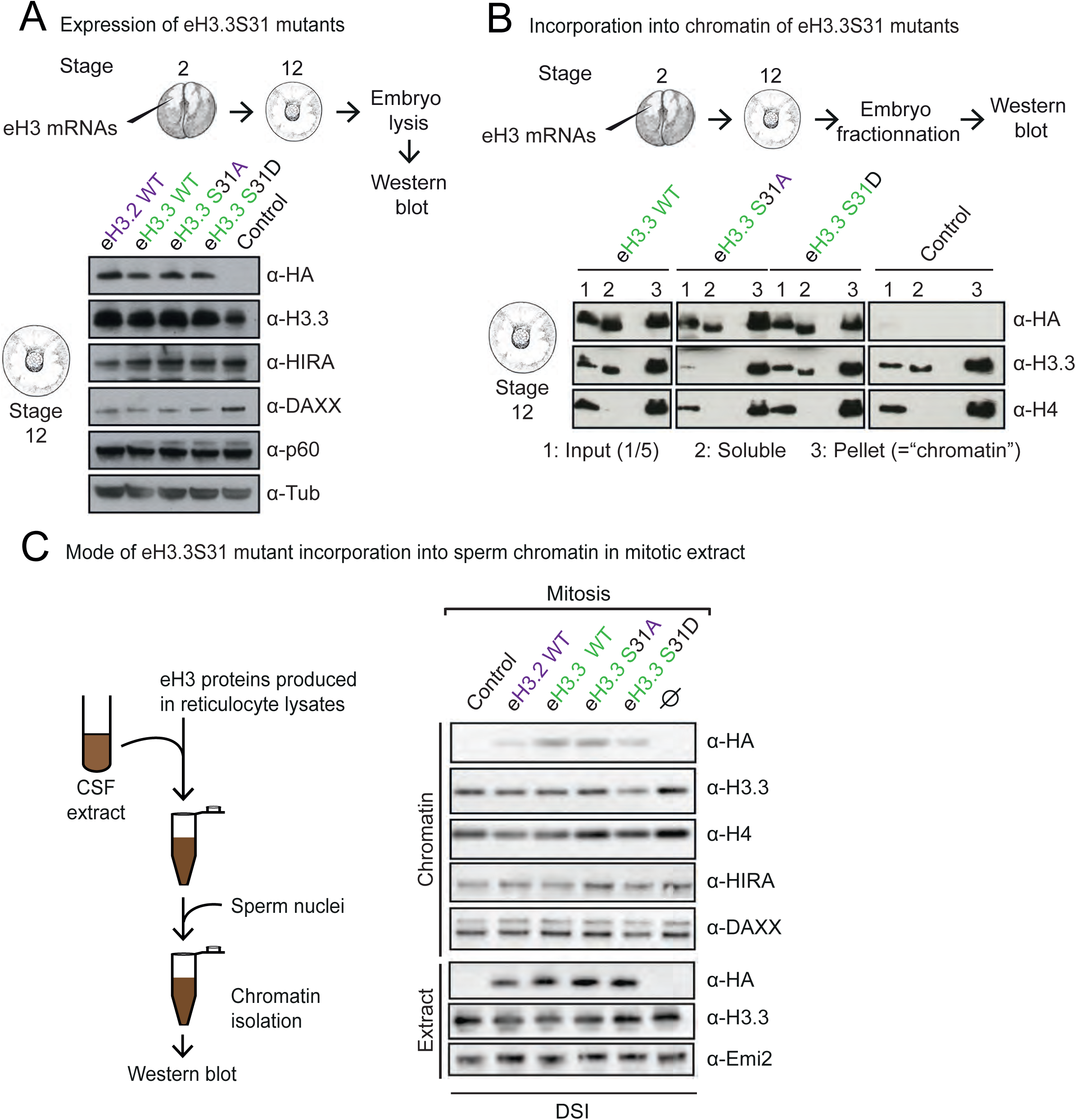
Mutant forms of H3.3S31 are all expressed and incorporated in a similar fashion. **A)** Western blot to control eH3.3S31 mutant expression in embryos. eH3.3S31 mutant expression compares with the other eH3 conditions in whole embryo extracts at stage 12. **B)** Western blots to control eH3.3S31 mutant incorporation in embryos. Fractionation of stage 12 embryos shows that these mutant remains insoluble at 600nm NaCl, and therefore most likely incorporated into chromatin. Note that both eH3.3 WT and Control conditions are reused in Supplemental Figure 2 B and 3 B since each condition used comes from the same western blot. **C)** Incorporation of eH3.3S31 mutant forms into sperm chromatin in mitotic extract. Purified nuclei remodeled in the CSF-arrested extracts supplemented with eH3.3 mutants were analyzed by WB with indicated antibodies. In mitotic extract where the only pathway available is the DSI, mutations of the residue 31 do not change the mode of incorporation of these forms.

